# Causes and consequences of nuclear gene positioning

**DOI:** 10.1101/118711

**Authors:** Sigal Shachar, Tom Misteli

## Abstract

The eukaryotic genome is organized in a manner that allows folding of the genetic material in the confined space of the cell nucleus, while at the same time enabling its physiological function. A major principle of spatial genome organization is the non-random position of genomic loci relative to other loci and to nuclear bodies. The mechanisms that determine the spatial position of a locus, and how position affects function, are just beginning to be characterized. Initial results suggest multiple, gene-specific mechanisms and the involvement of a wide range of cellular machineries. In this Commentary, we review recent findings from candidate approaches and unbiased screening methods that provide initial insight into the cellular mechanisms of positioning and their functional consequences. We highlight several specific mechanisms including tethering to the nuclear periphery, passage through replication and histone modifications that contribute to gene positioning in yeast, plants and mammals.

## Introduction

The genome within the nucleus of eukaryotic cells is organized in a complex, hierarchical manner that allows folding of DNA in a confined space and at the same time enables proper function, ensuring expression of the correct gene programs in the right place and at the right time (Box 1). The folding of DNA into a chromatin fiber is essential for the extensive compaction of the genome but is also critical for the functional regulation of genomes. The chromatin fiber is folded and looped in such a way that allows it to physically associate with other chromatin regions both in *cis* and in *trans* or with nuclear structures, creating interactions which may be functionally relevant (Cavalli and Misteli, 2013). As cells change their behavior and function, for example, during differentiation or malignant transformation, the physical interaction network is thought to be remodeled to reflect the functional status of the cell (Wang and Dostie, 2016).

A fundamental feature of genome organization is its non-random spatial organization. Individual genes and genome regions can assume different positions in the 3-dimensional space of the nucleus (Pombo and Dillon, 2015). In some cases, the position of a locus correlates with function (Pombo and Dillon, 2015; Volpi et al., 2000; Williams et al., 2002). For example, in mouse erythroid precursor cells and B cells, the β-globin gene when active colocalizes at discrete transcription hubs together with other active genes located at distal sites on the same chromosome (Eskiw et al., 2010). During differentiation, genes may be placed in specific compartments which determine their transcriptional status and affect cell identity and behavior. For example, during differentiation of olfactory sensory neurons, only one receptor is chosen to be expressed and comes to reside in an active compartment, whereas the other silent receptor genes aggregate in a spatially separated inactive compartment (Clowney et al., 2012). Of pathological relevance, in cancer tissues from breast and prostate, several genes change their nuclear position compared to normal non-cancerous tissues from the same individual, although these repositioning events do not necessarily correlate with gene activity (Leshner et al., 2016; Meaburn et al., 2009).

The non-random positioning of genomic loci is a fundamental property of genomes, yet, the molecular factors that control the positioning of genes in 3D are only just beginning to be uncovered. Two general approaches are currently used to identify factors that organize the interphase nucleus: candidate approaches, in which specific cellular factors are tested for their ability to affect the position of individual genes and, alternatively, unbiased screening approaches to identify novel players and pathways. Here, we review recent approaches and results that have shed light on this fundamental question in cell biology.

## Studying specific genome organizing components

The notion of non-random positioning of genes emerged from the empirical observation of preferential location of genes of interest when analyzed by fluorescence in situ hybridization (FISH) (Cremer et al., 2006). The particular position of a gene, for example near the nuclear pore or associated with heterochromatin, and the nature of the gene, such as its transcriptional activity or ability to be induced, often generated specific hypotheses for potential molecular mechanisms of positioning. The advantage of such candidate approaches is the clearly defined hypothesis, which allows precise experimental investigation of a potential mechanism. Since the nuclear periphery is one of the most prominent spatial features of the eukaryotic cell nucleus, it is often used as a reference point when assessing gene positioning and much of what we know about positioning factors relates to the proximity of genes to the nuclear envelope. The molecular mechanisms that control activation or repression of genes upon repositioning to the nuclear periphery have been studied in several model organisms (Fig. 1).

**Fig. 1.**
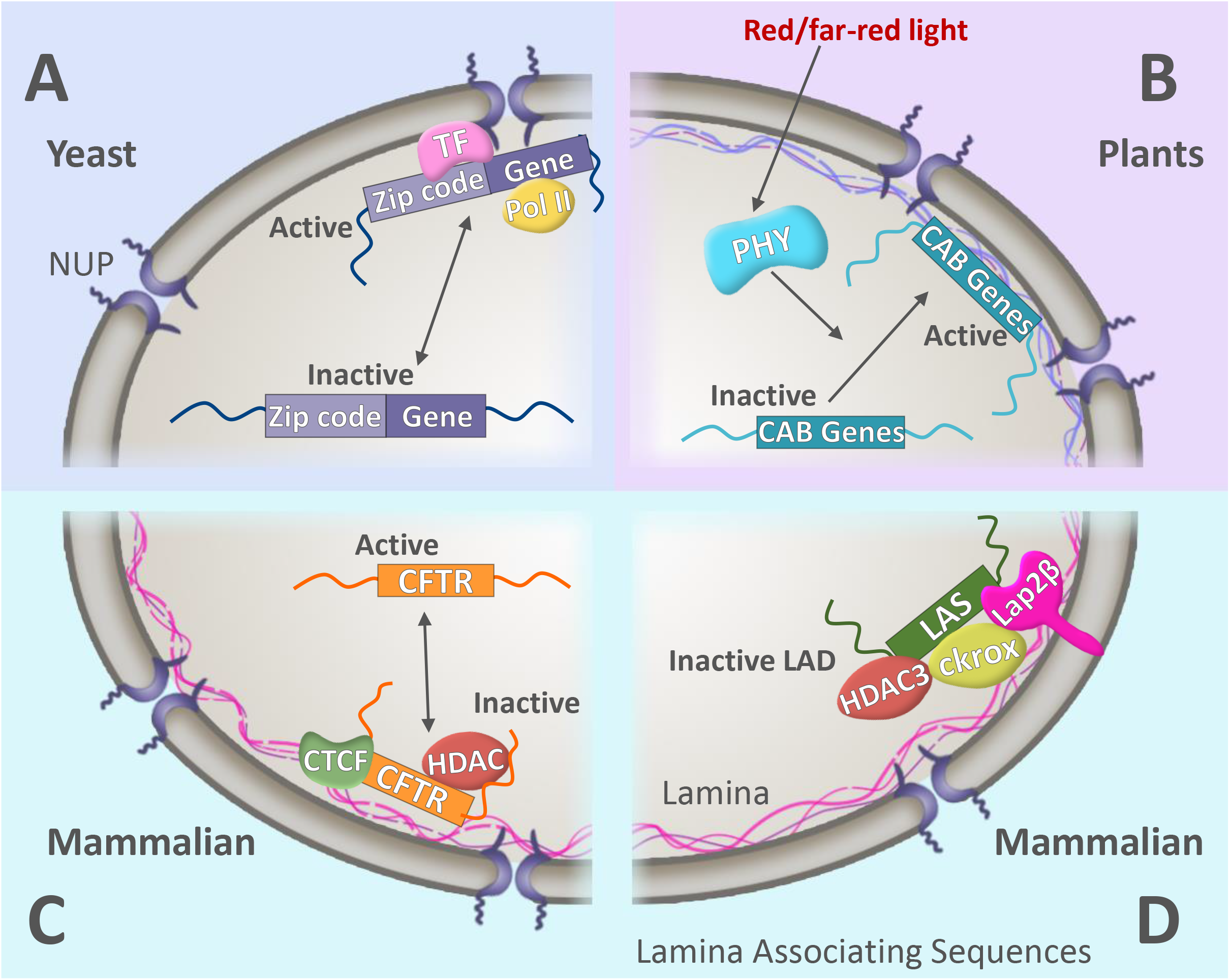
mechanism for positioning to the nuclear periphery in various organisms. Movement of a genomic locus towards the periphery of the nucleus has functional implications on its transcriptional activity in different organisms. (A) In yeast, specific zip-code sequences that reside in the promoter region of many genes interact with a transcription factor (TF) and with a nuclear pore complex (NUP) protein, leading to its re-localization to the periphery along with an increase in expression. (B) Gene activation and localization in response to light in plants. Following exposure of *Arabidopsis thaliana* cells to red or far-red light, activation of photoreceptors (PHY) leads to re-positioning of the CAB gene locus to the nuclear periphery where CAB genes are activated. (C) In mammalian cells, repression of the CFTR gene locus correlates with its peripheral localization, which occurs through a mechanism that involves CTCF, lamins and a histone deacetylase. (D) The mechanisms for anchoring lamin-associated domains (LADs) to the nuclear periphery in mammalian cells involve specific lamina associating sequences (LASs), the nuclear factors Lap2β, histone deacetylase 3 (HDAC3) and the transcription factor cKrox.

In yeast, several inducible genes are targeted to the nuclear periphery upon their activation (Ahmed et al., 2010; Brickner et al., 2007; Brickner and Walter, 2004). Recruitment of these genes to the nuclear periphery is mediated by their physical interaction with the nuclear pore complex (NPC) and constrains movement of the gene (Ahmed et al., 2010; Brickner et al., 2007; Brickner and Walter, 2004). Specific sequences within the promoter region of several yeast genes *(INO1, GAL1, HSP104,* and *TSA2*) are necessary for targeting these genes to the periphery and for interaction with NPCs. These sequences function as DNA zip codes and are sufficient to target any locus to the NPC when they are artificially introduced into a different genomic region. Additionally, endogenous genes with the same DNA zip code sequences cluster in space and are separated from other genes that contain a different zip code sequence in their promoter (Brickner et al., 2012) (Fig. 1A).

These peripheral targeting DNA zip codes comprise of a core 6-20-bp sequence which is distinct from the known upstream promoter elements (Ahmed et al., 2010). They appear hundreds of times in the yeast genome and are enriched in promoters of NPC-interacting genes or stress-induced genes. The zip codes use several strategies to determine the position of genes, but all require association between a zip code sequence, a transcription factor (TF) that binds the zip code and a NPC protein. One example is the *INO1* gene promoter, which is targeted to the nuclear periphery by the Put3 transcription factor that binds the DNA zip code. The recruitment of the Put3 TF and its binding to the specific zip code in the *INO1* promoter is regulated by the histone deacetylase complex Rpd3(L) and leads to increased transcription (Randise-Hinchliff et al., 2016). The peripheral re-positioning of other genes such as *PRM1* is regulated through a different mechanism and mediated by the Ste12 TF, which is constitutively bound to the promoter region and is repressed by the inhibitor Dig2. Upon stimulation, the MAPK pathway leads to phosphorylation of Dig2, which removes the inhibition of Ste12, thereby leading to positioning of the gene to the NPC and its activation (Randise-Hinchliff et al., 2016).

Similar to yeast, activation of genes upon their recruitment to the nuclear periphery also occurs in plant cells. In *Arabidopsis,* the chlorophyll a/b-binding protein (CAB) gene locus, which contains a 7kb cluster of three members of the *CAB* gene family (*CAB1-3*) on chromosome 1, re-positions to the nuclear periphery in response to red or far-red light (Feng et al., 2014) (Fig. 1B). In the dark, these genes are repressed and physically retained in the nuclear interior of mesophyll cells by a number of master repressors of photoreceptor signaling (DET1, COP1 and PIFs). The rapid repositioning of the locus to the periphery upon light exposure is mediated by the red and far-red photoreceptors phytochromes (PHYs) and their signaling components, which are the main components in *Arabidopsis* responsible for sensing continuous monochromatic light. Interestingly, elevated transcription of the *CAB* genes was evident a few hours after re-positioning to the periphery, suggesting an additional regulatory step possibly by other factors, leading to positioning and activation (Feng et al., 2014).

In mammalian cells, a prominent gene cluster that is localized to the nuclear periphery is the cystic fibrosis transmembrane conductance regulator (CFTR) region on human chromosome 7q31 that contains three adjacent genes: GASZ, CFTR, and CORTBP2. When these three genes are transcriptionally inactive in neuroblastoma cells, they are localized to the nuclear periphery and are associated with heterochromatin and the LAP2β protein (Zink et al., 2004) (Fig. 1C). In contrast, when actively transcribed in adenocarcinoma cells, they occupy an interior position and are associated with euchromatin. This gene-cluster has a differential spatial organization in various cell types, depending on the activity levels of each gene, with the actively transcribed gene sequestered from the silent genes and in a more internal position (Zink et al., 2004). Interestingly, in adenocarcinoma cells treated with the transcription inhibitor DRB, the CFTR gene relocalizes to the nuclear periphery concomitantly with a 50% reduction in its transcription. However, the transcriptional status of CFTR is not altered by treating cells with an inhibitor of histone de-acetylation (HDAC) despite the HDAC-induced change in nuclear positioning. These results suggest that transcriptional activity of these genes does not depend solely on their position, but also on their chromatin environment, histone modifications and interactions with other DNA elements. With regards to the molecular mechanism, CTCF, lamin A/C and active HDAC were found to contribute to the regulation of positioning (Muck et al., 2012) (Fig. 1C). For instance, knockdown of CTCF or laminA/C using siRNA in human cells led to a significantly more interior position of CFTR, but did not affect the position of the neighboring ASZ1 and CTTNBP2 genes. Treating Calu-3 adenocarcinoma cells with the HDAC inhibitor TSA led to a similar outcome, whereby the CFTR promoter was highly enriched in histone H3Ac and H4Ac and was relocated to the nucleus interior (Muck et al., 2012).

The CFTR locus is not the only gene cluster to associate with the nuclear lamina in its inactive state and release upon activation. The same phenomenon occurs in mouse cells for the β-globin and IgH loci (Kosak et al., 2002; Ragoczy et al., 2006) and in *C. elegans* for several developmentally regulated genes (Meister et al., 2010). The positioning mechanisms appear to rely on the interaction of the inner nuclear membrane (INM) and the underlying lamina with chromatin regions dispersed throughout the linear genome, which are referred to as lamina associating domains (LADs) (Guelen et al., 2008). One mechanism by which the INM-lamina compartmentalize chromatin domains and silence genes is through specific DNA sequences within LADs. These sequences termed lamina-associating sequences (LASs) were found at the *IgH* locus and the Cyp3a gene cluster that comprise a continuous LAD on chromosome 12 and 5, respectively, in mouse cells. These sequences are sufficient to target chromatin to the lamina upon their integration into an ectopic site and colocalize with lamin B at the end of anaphase when the lamina begins to re-form around chromatin, thereby directing the peripheral localization of LADs (Zullo et al., 2012) (Fig. 1D). Specific DNA motifs in LASs bind the transcriptional repressor cKrox, which in turn interacts with HDAC3 and Lap2β to direct association with the lamina and silencing of transcription, likely acting in concert with additional INM proteins. In support, knockdown of any of these factors reduces the association and increases transcriptional activity of a reporter gene and of endogenous murine LAD genes (Zullo et al., 2012). As cKrox associates with mitotic chromosomes it may serve as a docking point for additional proteins, such as lamins and Lap2β, during early G1 phase when cells reorganize their chromatin and so might facilitate nuclear envelope assembly and genome organization in parallel.

Lamin A/C and the lamin B receptor (LBR), which are both peripheral NE-binding proteins, have further been implicated in chromatin organization and accurate gene positioning based on observations in mice (Solovei et al., 2009). While in most mammalian cells heterochromatin is enriched at the periphery of the nucleus and relies on the integrity of the nuclear lamina and associated proteins, in striking contrast, in rod photoreceptor cells of nocturnal mammals, compact, silent heterochromatin resides in the center of the nucleus (Solovei et al., 2009). This inversion of chromatin organization depends on low expression of lamin A/C and LBR, as cells with conventional chromatin pattern express high levels of these proteins, whereas the inverted rod cells do not (Solovei et al., 2013). Indeed, postmitotic cells from mice, in which both lamin A/C and LBR have been knocked out, show inversion of chromatin in various cell types, confirming the necessity of these proteins to maintain peripheral heterochromatin. Conversely, ectopic expression of these factors in rod cells prevents inversion and leads to conventional chromatin architecture. LBR and lamin A/C are expressed at different developmental stages, which is consistent with the opposite effects their loss has on the transcription of muscle-related genes in myoblasts; here loss of LBR increases their expression, whereas loss of lamin A/C decreases expression (Solovei et al., 2013).

LBR has also been implicated in the positioning of olfactory receptor (OR) gene clusters in olfactory sensory neurons (OSNs); out of the 2,800 OR alleles, only a single allele is chosen to be expressed in an individual cell (Buck and Axel, 1991; Imai et al., 2010). The expressed allele harbors distinctive chromatin structure and histone modifications compared to the other, inactive ORs, which cluster together in 3D space and colocalize with known heterochromatic marks, such as H3K9me3, H4K20me3 and HP1β (Clowney et al., 2012; Magklara et al., 2011). In contrast, the single active OR allele is located in a euchromatic domain, which is enriched in Pol II or H3K27Ac and does not show overlap with any of the heterochromatin markers (Clowney et al., 2012). Similar to rod cells, LBR is absent from OSNs and has a role in the aggregation of inactive ORs as indicated by the fact that restoration of LBR expression in OSNs leads to a disruption in the formation of inactive OR foci, decondensation of heterochromatin and misregulation of OR expression, despite the fact that heterochromatin histone marks, such as H3K9me3 and H4K20me3, are retained (Clowney et al., 2012).

Despite the fact that in many human cell types, the NE is mostly lined by heterochromatin, genomic regions immediately at NPCs are generally more open and devoid of heterochromatic marks (Capelson and Hetzer, 2009). Genomic regions that interact with NPC proteins such as Nup153 and Nup93 are mainly comprised of gene enhancers and are closer to transcription start sites (Ibarra et al., 2016). Moreover, knock-down of Nup153 results in a change in expression of genes associated with enhancers that bind NPCs, suggesting that NPC proteins regulate the transcriptional activity of these genes in a selective manner (Ibarra et al., 2016). The importance of NPC proteins in controlling gene expression is also demonstrated by mutations in NPC components that alter the expression of genes and lead to a variety of tissue pathologies in flies and human (Capelson and Hetzer, 2009; Nofrini et al., 2016).

Taken together, these studies point to a role for several nuclear lamina and NE-proteins in organizing heterochromatin and tethering it either to the nuclear periphery or to distinct foci, while maintaining high compaction and low expression levels. It is likely that additional NE-proteins, which remain to be identified, participate in this anchoring and silencing process. In addition, in mammalian cells, these candidate factors are involved in the regulation of silencing at the periphery. However, additional factors that mediate the peripheral location of active regions likely exist as well.

## Identifying gene positioning factors using unbiased screens

While candidate approaches are powerful in elucidating a specific positioning mechanism, they are unable to identify unanticipated pathways and it is often difficult to establish the global relevance of factors identified using these approaches. A powerful alternative approach to discover and characterize factors and pathways that spatially organize the genome are genetic or RNA interference (RNAi)-based screens. These methods represent a true discovery approach and have the advantage of not making any assumptions about involved pathways. Screening strategies to identify gene positioning factors have recently become feasible due to the development of high-throughput FISH methods using high-content imaging (Joyce et al., 2012; Shachar et al., 2015a; Shachar et al., 2015b) in which the position of a large number of genes or large number of conditions, such as RNAi knockdowns, can be probed in a single experiment. These approaches have yielded the most comprehensive sets of gene positioning factors and have led to the characterization of several novel positioning pathways.

In the worm *C. elegans* and in flies, lamins and lamin-binding proteins are required for silencing of specific heterochromatin-associated genes (Mattout et al., 2011; Shevelyov et al., 2009), but the factors involved in lamina-attachment and silencing and their mechanism of action were not fully understood. To address this question, a genome-wide RNAi screen was performed in *C. elegans* embryos using stably integrated repetitive arrays, which are located at the nuclear periphery and are silenced (Towbin et al., 2012). By monitoring both increase in expression of an integrated GFP reporter gene and detachment from the nuclear periphery, two cellular factors were identified as being important for both tethering and silencing at the periphery: S-adenosyl methionine synthetase (SAMS)-3 and SAMS-4, two enzymes that generate S-adenosylmethionine (SAM), a universal methyl group donor for many DNA and protein methylation reactions in eukaryotes. In the absence of these enzymes, there is a massive reduction in histone H3 methylation specifically on K9, K27 and K36 throughout the nucleus. This methylation is performed by the methyltransferases MET-2 and SET-25 (homologous to SETDB1 and Suv39h1/2 / EHMT1/G9a in mammals, respectively) and they both act redundantly to anchor heterochromatin at the nuclear periphery. When both methyltransferases are lost, the silenced array is completely released from the periphery, its interaction with the lamina is reduced and expression of the reporter gene is increased, suggesting that H3K9 methylation is sufficient to anchor the array and tether heterochromatin (Towbin et al., 2012). To characterize the particular nuclear factor that directly anchors methylated H3 and is physically responsible for tethering at the periphery, an additional RNAi screen in *C. elegans* searched for factors that anchor a heterochromatic reporter gene to the periphery (Gonzalez-Sandoval et al., 2015). This screen identified a single uncharacterized chromodomain factor CEC-4 that is responsible to anchor heterochromatin to the periphery in a manner similar to double knock-down of *met-2* and *set-25* methyltransferases, but did not alter expression levels of the reporter gene. CEC-4 localizes to the nuclear periphery in cells from all embryonic stages independent of H3K9 methylation and of lamins and binds to H3K9me3 through its chromodomain. CEC-4 affects chromatin organization and distribution of endogenous chromosomes and loss of CEC-4 leads to detachment of specific chromosomal sequences from the periphery. Loss of H3K9 methylation by downregulating methyltransferases, upregulates many genes in embryos; however, the loss of CEC-4 led to robust upregulation of a single gene in the *C. elegans* genome, demonstrating that CEC-4 unlike H3K9 methylation, serves mostly to position chromatin to the periphery and not to silence genes in general (Gonzalez-Sandoval et al., 2015). Peripheral anchoring by CEC-4 helps to stabilize tissue-specific differentiation programs. When *cec-4* is mutated in muscle-progenitor cells, these cells can continue their development into other tissues as well. These RNAi screens in worms identified H3K9 methylation and CEC-4 as essential in anchoring heterochromatin to the inner membrane (Gonzalez-Sandoval et al., 2015; Towbin et al., 2012). Interestingly, proper heterochromatin organization was also shown to be compromised in mammalian cells with disrupted H3K9 methylation (Kind et al., 2013; Pinheiro et al., 2012); however, no functional homolog of CEC-4 was discovered in higher organisms and it remains to be determined how globally applicable this tethering mechanism is.

Heterochromatin tethering to the INM has been shown to be important for chromatin organization, but it does not address other types of non-random gene positioning. One functionally highly relevant occurrence of gene positioning is pairing of homologous chromosomes, which can occur in somatic tissues and affects gene regulation and DNA repair in the fly *Drosophila melanogaster* (Henikoff and Comai, 1998; Kassis, 2002; Rong and Golic, 2003). Using high-throughput FISH and a whole-genome RNAi screen in *Drosophila* cell culture,Joyce et al. identified cellular factors that either promote or antagonize somatic pairing of endogenous centromeric heterochromatic loci (Joyce et al., 2012). In total, 40 candidate pairing-promoting genes and 65 anti-pairing factors were uncovered, most of them not previously implicated in gene positioning. Many of these factors contribute to the normal progression of cell cycle, such as nucleotide biosynthesis, G1/S transition factors and replication Joyce et al., 2012). Some, such as the SCF E3 ubiquitin ligase complex and the anaphase promoting complex APC, promote pairing of homologous chromosomes in both heterochromatic and euchromatic regions, suggesting a more global role in genome organization. Intriguingly, a few of the discovered factors that counteract pairing of homologous chromosomes were proteins that can lead to chromatin compaction, including HP1a, ORC1 and SLE, as well as components of the condensin II complex, such as CAP-H2, CAP-D3 and SMC2. Prevention of pairing of heterochromatin between homologues may be beneficial to limit crosstalk between chromosomes that can lead to errors in replication and repair, since repetitive heterochromatic regions pose a challenge to replication forks as they tend to stall at repetitive sequences and sometimes collapse, which can lead to loss of heterozygosity and, in severe cases, to the formation of chromosome bridges during mitosis (Mirkin, 2006). Interestingly, the observed involvement of replication and cell cycle regulators in both pairing and anti-pairing (Joyce et al., 2012) also suggests that faithful progression through the cell cycle is crucial for genome organization - a concept which also emerged from our recent gene positioning screen in human cells (Shachar et al., 2015b).

Identification of factors that determine the position of endogenous genes or single loci have been limited by the difficulty in detecting and measuring accurately and reliably the nuclear position of genes in a large number of samples. A novel method we termed HIPMap (high-throughput imaging positioning mapping), in which automated high-throughput FISH and image analysis are used to determine the position of multiple loci, allowed us to overcome the limitation of monitoring artificial arrays or repetitive sequences and to probe positioning of endogenous genes with variable expression patterns and chromatin environments (Shachar et al., 2015b). We used HIPMap in an unbiased RNAi screen in human cells to monitor changes in positioning of four endogenous loci with distinct transcriptional features and nuclear locations, including active or silent genes and peripheral or internal loci. The screen in human skin fibroblasts uncovered 50 cellular factors that are required for proper gene positioning, most of them not previously implicated in nuclear organization or gene positioning. Notably, most of the factors identified affected the positioning of some of the target loci but not all, indicating that gene positioning is not determined by a single dedicated machinery, but rather that multiple pathways and mechanisms contribute to position various loci and to organize compartments with diverse expression status and chromatin state (Shachar et al., 2015b). By monitoring expression levels of several genes following re-positioning it was shown that gene-positioning is not tightly linked with activity, demonstrating that positioning and expression can be uncoupled, which is in line with previous findings (Therizols et al., 2014).

One prominent group of cellular factors identified in our HIPMap screen were DNA replication proteins, which is in agreement with the screen conducted in *Drosophila* cells (Joyce et al., 2012). Interference with several replication factors through RNAi resulted in re-position of the target loci. Importantly, when replication was slowed down by drug treatment, several endogenous loci mis-localized, suggesting that the process of replication itself, rather than just the replication factors, contributes to gene positioning (Shachar et al., 2015b). It is tempting to speculate that the epigenetic code and histone modifications that are re-assembled during S phase contribute to positioning and serve as nucleation sites for the organization of the nucleus in the subsequent G1 phase (Fig. 2). The finding that chromatin domains and TADs are lost during mitosis and get re-established in early G1 (Naumova et al., 2013) may suggest a scenario in which immediately after the daughter strand is replicated, new modified histones are assembled onto the DNA, followed by mitosis in which specific histone modifications maintain interaction with nuclear proteins as was demonstrated for laminB and LAD-derived sequences that colocalize at the end of anaphase (Zullo et al., 2012). Following mitosis, when nuclei re-organize their chromatin, these interactions then serve as docking points for further assembly of similarly modified chromatin and, in this way, determine the position of genomic loci (Fig. 2). Live-cell imaging experiments that can track the position of an endogenous locus and its interactions with several proteins throughout the cell cycle will be required to confirm this hypothesis.

**Fig. 2.**
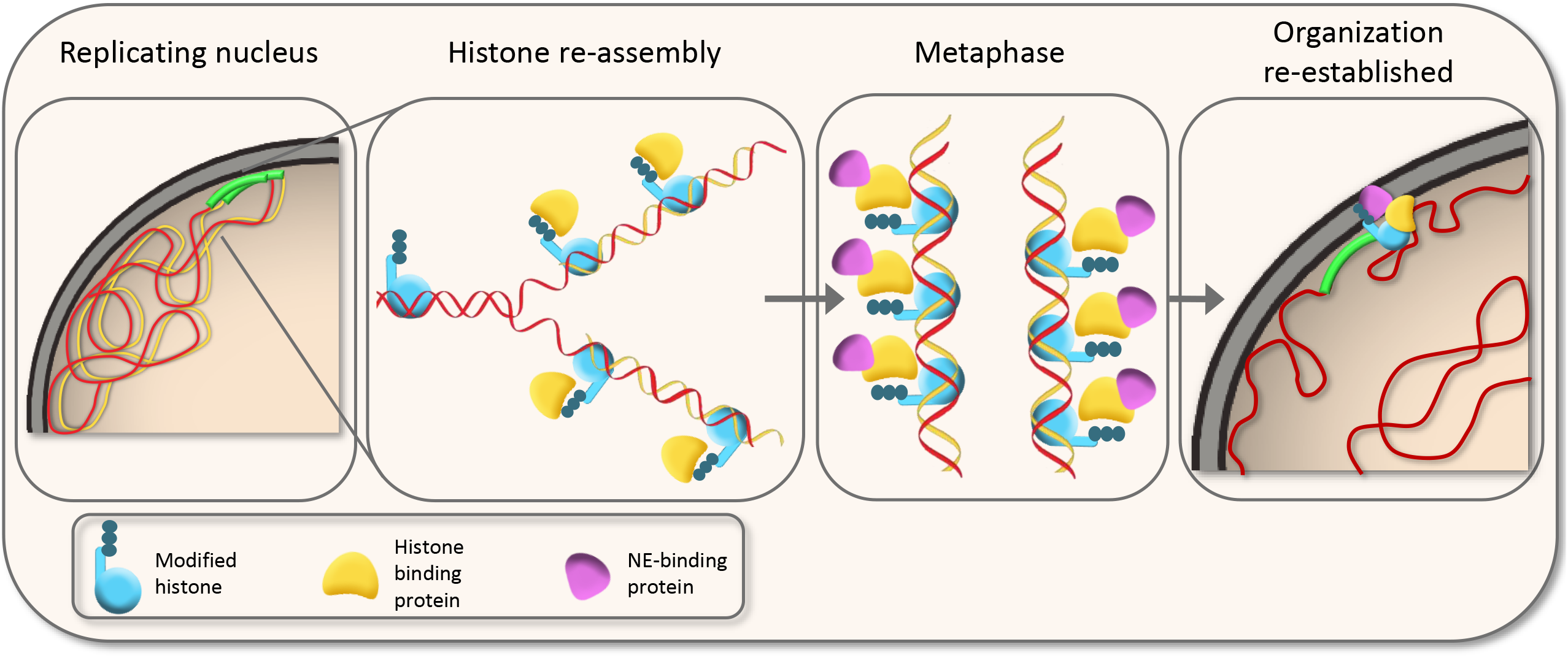
A putative mechanism for positioning by replication. Genomic loci have preferred nuclear positions that are re-established following chromosome condensation and breakdown of the nuclear envelope during mitosis. During replication, specific histone modifications contribute to the binding of additional proteins which stay attached to chromatin throughout mitosis. When cells re-organize their chromatin in early G1, these proteins in turn bind scaffold proteins, such as the lamina or nuclear envelope (NE)- proteins, ensuring accurate positioning of the locus.

In addition to replication, chromatin-remodeling complexes also appear to contribute to positioning. Three identified hits from the HIPMap screen, ARID1A, SMARCD2 and SMARCD3, are a part of the BAF complex (BRG1 associated factors), an ATP-dependent chromatin remodeler complex (SWI/SNF) important for controlling cell fate and lineage specification (Ho and Crabtree, 2010; Reisman et al., 2009; Wilson and Roberts, 2011). In human cells, the BAF complex affects positioning of peripheral low-expressed genes and leads to re-positioning of these loci to a more peripheral location (Shachar et al., 2015b). Deletion of several subunits of this complex in mouse embryonic fibroblasts led to a reduction in nucleosome occupancy at transcription start sites and up to 2kb up-and downstream with only a mild effect on transcription (Tolstorukov et al., 2013). Taken together, it is possible that the BAF complex contributes to positioning of gene loci by altering the higher-order chromatin structure and in this way promoting local contacts between similar chromatin domains. This mechanism could be specific to genes or transcribed regions, since a control locus within a gene-less region was not affected by the loss of these factors (Shachar et al., 2015b).

Another mechanism shown to alter positioning is methylation of H3K9, similar to the situation in *C. elegans* (Towbin et al., 2012). The H3K9 methyltransferase SETDB2 and the H3K27 demethylase KDM6A both contribute to accurate positioning of several endogenous loci (Shachar et al., 2015b). Similar to worms, it appears that in humans, methylation patterns of histone H3 are important in tethering genomic regions to the periphery, thereby sequestering them from actively transcribed regions and keeping them silent. Additional experiments are required to probe the exact mechanism by which human cells retain and silence chromatin at the periphery.

## Gene positioning and gene function

The obvious and long-standing question, of course, is whether the position of a gene affects its activity. The position of a gene can be measured relative to the periphery of the nucleus or relative to another locus or nuclear compartment, such as nucleolus or transcription hubs. Anecdotal examples from several organisms and biological systems show that expression patterns of genes change when their nuclear position changes. One such example is the epidermal differentiation complex in keratinocytes that is frequently positioned outside of its chromosome territory when active compared to lymphoblasts, in which it acquires a more internal position concomitant with gene silencing (Williams et al., 2002). Another example is the major histocompatibility complex locus on chromosome 6 which loops outside of its chromosome territory more frequently in response to gene activation by interferon-gamma (Volpi et al., 2000). Furthermore, gene-rich loci generally cluster and are separated from gene-poor areas that tend to aggregate near the periphery or in peri-centromeric heterochromatin in mammalian cells (Pombo and Dillon, 2015). Additionally, as discussed above, CFTR moves away from the periphery upon activation, whereas the yeast genes *INO1, GAL1, HSP104,* and *TSA2* are moving to the periphery to associate with NPCs upon activation (Ahmed et al., 2010; Brickner and Walter, 2004; Muck et al., 2012; Zink et al., 2004). In contrast, some loci change position with no effect on their activity, for example in human mammary cells, there was no correlation between locus positioning and gene expression for several tested loci (Meaburn and Misteli, 2008). Similar results were obtained in human skin fibroblast where changes in positioning were uncoupled from transcription for some genes (Shachar et al., 2015b). This is in line with the finding that local decondensation of chromatin is sufficient to reposition an endogenous locus, whereas transcriptional activation had no effect on positioning (Therizols et al., 2014).

Several genes change their expression as they associate or dissociate from the periphery (Ahmed et al., 2010; Zink et al., 2004; Zullo et al., 2012). To directly test whether association of a locus to the periphery affects gene activity, several studies tethered a genomic locus to the periphery using a chimeric protein that binds both a specific DNA sequence and a peripherally localized protein. In one approach, an artificial target sequence containing repeats of the *E. coli* Lac Operator (LacO) was tethered to the nuclear periphery by using a chimeric protein lacI-mCherry - lamin B1 (Kumaran and Spector, 2008). Tethering of the locus to the periphery required passage through mitosis and it stayed tethered for subsequent cell cycles. The transcription machinery was recruited to the tethered locus at the same efficiency as to a control locus and RNA transcripts were visible, indicating that tethering a genomic locus to lamin B is not sufficient to inhibit transcription under these conditions. A very similar assay scheme, albeit in mouse cells, used a reporter construct made of GFP-LacI-DEMD targeted to the inner nuclear membrane by a segment of the inner membrane protein emerin (Reddy et al., 2008). Consistent with other reports, this protein tethered the LacO array to the nuclear periphery in a process that required breakdown of the nuclear envelope through mitosis. This test gene was transcriptionally repressed as a consequence of repositioning to the inner membrane, which was accompanied by histone H4 de-acetylation of the tethered genomic locus. An additional fusion protein comprised of LacI-Lap2β was used to tether two endogenous loci on chromosomes 4 and 11 to the nuclear periphery in human cells leading to reduced expression of several endogenous genes located adjacent to the tethered region, but not affecting expression of other nearby genes that also altered their nuclear position (Finlan et al., 2008). Consistent with the notion that reduced expression of tethered genomic regions is due to histone H4 hypo-acetylation, the expression of the reporter gene and of endogenous genes was increased following treatment with the HDAC inhibitor TSA (Finlan et al., 2008). Interestingly, despite the contrasting results regarding the resulting expression patterns, all tethering experiments that used a fusion protein required passage through mitosis to establish the peripheral gene positioning. Possible explanations for how physical attachment of a mammalian gene to the nuclear periphery promotes its transcriptional repression include the assembly of repressive marks, such as methylation of histone H3 at Lysine 9, inclusion in a repressive environment where inner membrane proteins interact with chromatin modifying enzymes and transcription repressors, as well as physical sequestration from activating components such as RNA pol II centers.

## Concluding remarks

The position in the nuclear space is a basic property of every genomic locus. Characterization of higher-order organization of chromatin and of the mechanisms that determine spatial positioning of genes is vital for our understanding of genome function in health and disease. Over the past decade, efforts have been made to describe and characterize specific factors and mechanisms that determine genome organization in various biological contexts and organisms. The least understood aspects of spatial genome organization are the mechanisms that determine where a gene or genome region is localized in the cell nucleus. Traditional candidate approaches and, more recently, unbiased screening approaches have provided first insights. Several conclusions have emerged: despite some overlap between factors that organize the genome such as lamins, LBR and heterochromatin marks, it is clear that there is no single determinant or mechanism that is solely responsible to localize chromatin, but rather various mechanisms are in place to allow the diverse organization in different organisms and cell types. Moreover, different loci are organized by diverse mechanisms, ranging from physical tethering by lamin-binding proteins (Clowney et al., 2012; Solovei et al., 2013) to specific DNA sequences that direct interactions (Ahmed et al., 2010; Brickner et al., 2012; Zullo et al., 2012) and histone modifications that in turn interact with adaptor proteins (Gonzalez-Sandoval et al., 2015; Towbin et al., 2012).

Major open questions are how gene position is linked to gene activity, whether a change in positioning can lead to different expression levels and whether transcription is required for determining gene position. Numerous studies in the past few years show that in some cases transcription can be uncoupled from positioning and that some genes change their activity following re-positioning while others do not, again suggesting gene-specific behavior (Hakim et al., 2011; Kumaran and Spector, 2008; Meaburn and Misteli, 2008; Shachar et al., 2015b; Takizawa et al., 2008; Williams et al., 2006). The application, particularly in combination, of biochemical methods such as chromosome conformation capture methods and single-cell imaging methods may begin to address these issues in the near future. One intriguing notion is the possible use of gene positioning as a bio-marker. Indeed, it has been shown that some genes change their position in cancer cells when compared to normal cells from the same individual and that this phenomenon can be harnessed to serve as a diagnostic tool to identify cancerous cells in various disease stages (Leshner et al., 2016; Meaburn, 2016; Meaburn et al., 2009).

Nuclear and genome organization and the positioning of genes are key aspects of normal and healthy cell function and are linked to cellular nuclear functions, in particular to gene activity and cell fate. There is no doubt that by utilizing novel techniques and the knowledge we have gained from recent observations, we will be able to further characterize the factors that organize the genome in the 3dimensional space of the nucleus and to uncover how they are involved in normal cell function and disease.

### Box 1 Basic organization of chromatin in the nucleus

The genome is hierarchically organized starting with the nucleosome, consisting of an octamer of the four core histones H2A, H2B, H3 and H4, around which DNA is wrapped. Chromatin folds in higher order fibers and ultimately into so called topologically associating domains (TADs), which bring genomic regions into close spatial proximity to allow regulation such as promoter-enhancer association (Dekker and Mirny, 2016). The discrete nature of TADs appears functionally important for either preventing or allowing physical interactions between genes and their regulatory elements as disruption of TAD integrity leads to gene mis-expression (de Wit et al., 2015; Guo et al., 2015; Lupianez et al., 2015). TADs themselves further associate with each other to form nuclear compartments that differ in histone modifications, density and nuclear position and may either be transcriptionally active or silent (Imakaev et al., 2012; Lieberman-Aiden et al., 2009). Functionally distinct compartments also differ in their association with nuclear bodies, with silent, compact heterochromatin associating with the nuclear lamina or the nucleolus, whereas open euchromatin associates with transcriptional hubs containing foci enriched in active RNA polymerase II; these are thought to be generated by non-random inter-and intragenic interactions between distal gene regions (Eskiw et al., 2010). Several structural proteins such as CCCTC-binding factor (CTCF), cohesin and Mediator have been implicated in organizing the basic units of chromatin, such as loops and TADs (Dekker and Mirny, 2016), and the boundaries of TADs contain binding sites for CTCF in opposite orientation which have been proposed to interact through loop extrusion activity of cohesins to form an insulated TAD (Bouwman and de Laat, 2015; Goloborodko et al., 2016; Rao et al., 2014; Sanborn et al., 2015).

